# Cohesin activity accelerates the homology search

**DOI:** 10.64898/2026.06.08.730890

**Authors:** Tylar Matsuo, Alberto Marin-Gonzalez, Taekjip Ha

## Abstract

Homologous recombination (HR) repair preserves genome integrity in post-replicative cells by orchestrating templated repair of DNA double-strand breaks (DSBs). A critical stage of HR is the homology search, the process by which the DSB is brought in contact with a repair template, which is often the replicated locus. Using molecular dynamics simulations, we model the mammalian homology search, with focus on the role of the chromatin architecture factor cohesin. Our simulations recapitulate key experimental findings, including the distribution of genomic loci interrogated by the DSB and the chromatin interactions that accompany the homology search. We show that cohesin-mediated loop extrusion greatly accelerates the search process, and this effect is further enhanced by anchoring loop-extruding cohesin at DSB sites and recruiting a cohesive cohesin clamp that stabilizes DSB-sister chromatid interactions. We reveal that cohesin’s contribution to accelerating the search scales linearly with TAD size and becomes more pronounced when breaks occur in large TADs. We also show that chromatin loops along the broken and the sister chromatid play different roles in the search: the former establish initial contact between the break site and sister chromatid, whereas the latter promote scanning along the sister chromatid. Our findings indicate that coordinated activity of loop-extruding and cohesive cohesin transforms the homology search from 3D diffusion into a fast 1D scanning process.

## INTRODUCTION

DNA double-strand breaks (DSBs) are among the most toxic forms of DNA damage, leading to mutations and chromosomal abnormalities associated with cancer when repaired incorrectly (1). Homologous recombination (HR) is a mechanism for DSB repair that operates in the G2 and S phases of the cell cycle (2). During HR, the DNA ends on both sides of the DSB get resected, exposing a 3’ single-stranded DNA overhang that is coated with the RAD51 recombinase, forming a nucleoprotein filament known as the presynaptic filament. Subsequently, the presynaptic filament “searches” for a homologous sequence to use as a template for repair in a process known as a homology search (2, 3). Once an appropriate template is found, HR proceeds via invasion of the donor template, strand synthesis, and resolution of repair intermediates.

In somatic cells, the sister chromatid provides a preferred substrate for HR repair compared to other chromosomes (4, 5). This preference is likely a result of sister chromatid cohesion, which is mediated by chromatid linkages established by cohesive cohesin, a form of the cohesin complex that keeps sister chromatids attached to each other from the time of DNA replication until mitosis (4–6). Consistently, early works in yeast and mammalian cells revealed that cohesin promotes sister chromatid recombination through its cohesion activity. Along these lines, a recent study reported that sororin, a key subunit of cohesive cohesin, is enriched at DSB sites undergoing HR and is required for the DSB to engage in chromatin contacts with the sister chromatid (7). In addition, recent works have revealed that cohesin also stimulates the homology search via formation of chromatin loops in the vicinity of the DSB site (7, 8). This involvement of loop-extruding cohesin is supported by Hi-C and ChIP-Seq experiments showing, respectively, cohesin-dependent interactions between the DSB and the broken chromatid and recruitment of loop-extruding cohesin to the broken chromatin at the time of the search (7, 8). The emerging model is that cohesin regulates the homology search via a dual mechanism: cohesive cohesin keeps the DSB in proximity to the sister chromatid and loop-extruding cohesin accelerates chromatin scanning (7, 8).

Yet, many questions remain open. What are the dynamics of this search and by how much does cohesin contribute? How do cohesive and loop-extruding cohesin coordinate with each other, and what are their relative contributions to the search? Does chromatin context, and specifically the positioning of topologically associating domain (TAD) boundaries, influence search kinetics?

Polymer simulations offer a prime tool to address these questions. Molecular dynamic simulations of chromosome folding identified loop extrusion as a mechanism for TAD formation (9), a model that has been backed by several experimental findings and is now well-accepted (10, 11). Simulations have also shown that cohesin can contribute to DSB end synapsis during non-homologous end joining repair (12), and have revealed how nucleolus formation and epigenomics-driven interactions shape 3D genome organization (13).

Here, we explore the role of cohesive and loop-extruding cohesin in the homology search using polymer molecular dynamics simulations. We build a model of sister chromatids within a replicated genome and simulate the homology search process in the presence of cohesive and loop-extruding cohesins. This model recapitulates several experimental findings, including the chromatin profile of the search (as given by RAD51 ChIP-Seq experiments), the effect of cohesin depletion on HR efficiency, and the chromosome conformation capture contacts associated with homology searches. Our simulations indicate that break-interacting cohesins reduce search times by approximately 2-fold. Furthermore, we find that TAD sizes influence search kinetics, with larger TADs leading to slower, more cohesin-dependent search processes. Finally, our computational model suggests a mechanism by which loop-extruding cohesins coordinate with cohesive cohesin and TAD boundaries to accelerate the homology identification process.

## RESULTS

### Loop extrusion accelerates search kinetics

We simulate chromatin as a polymer formed of monomers connected by harmonic bonds, each representing approximately 1 kb of DNA. Loop-extruding cohesins connect monomer positions to form topological loops and may load and unload randomly along the simulated polymer in accordance with processivity (defined as lifetime multiplied by the extrusion rate) and density (average separation) parameters. Loops grow as loop-extruding cohesin legs translocate in opposite directions along the polymer, drawing more chromatin into the enclosed loop (9, 14, 15). We assume DSB ends to be tethered in the main body of this paper unless otherwise stated, in line with experimental findings in yeast (16, 17).

We began with the simplified case of a search along one chromatid (Fig. 1A). We loaded loop-extruding cohesins onto a 150 Mb chromatin polymer with a processivity value of *λ*=120 and an average separation *d*=120, consistent with experimental and computational literature (9, 18). Similarly to (12, 14), we partitioned the 150 Mb polymer into 50 identical 3 Mb regions with a DSB centered in each, then averaged over each region and over each of 5 independent replicates of the simulation to achieve a total n=250 searches. We recorded the simulation timestep at which the DSB entered a 5 monomer-length threshold radius of a given monomer as a completed search to that monomer. We considered three different scenarios: simulations lacking loop-extruding cohesins, simulations with loop-extruding cohesins that do not interact with (i.e. translocate over) the DSB, and loop-extruding cohesins that become stabilized and anchored upon encountering the DSB (Fig. 1B).

**Figure 1:**
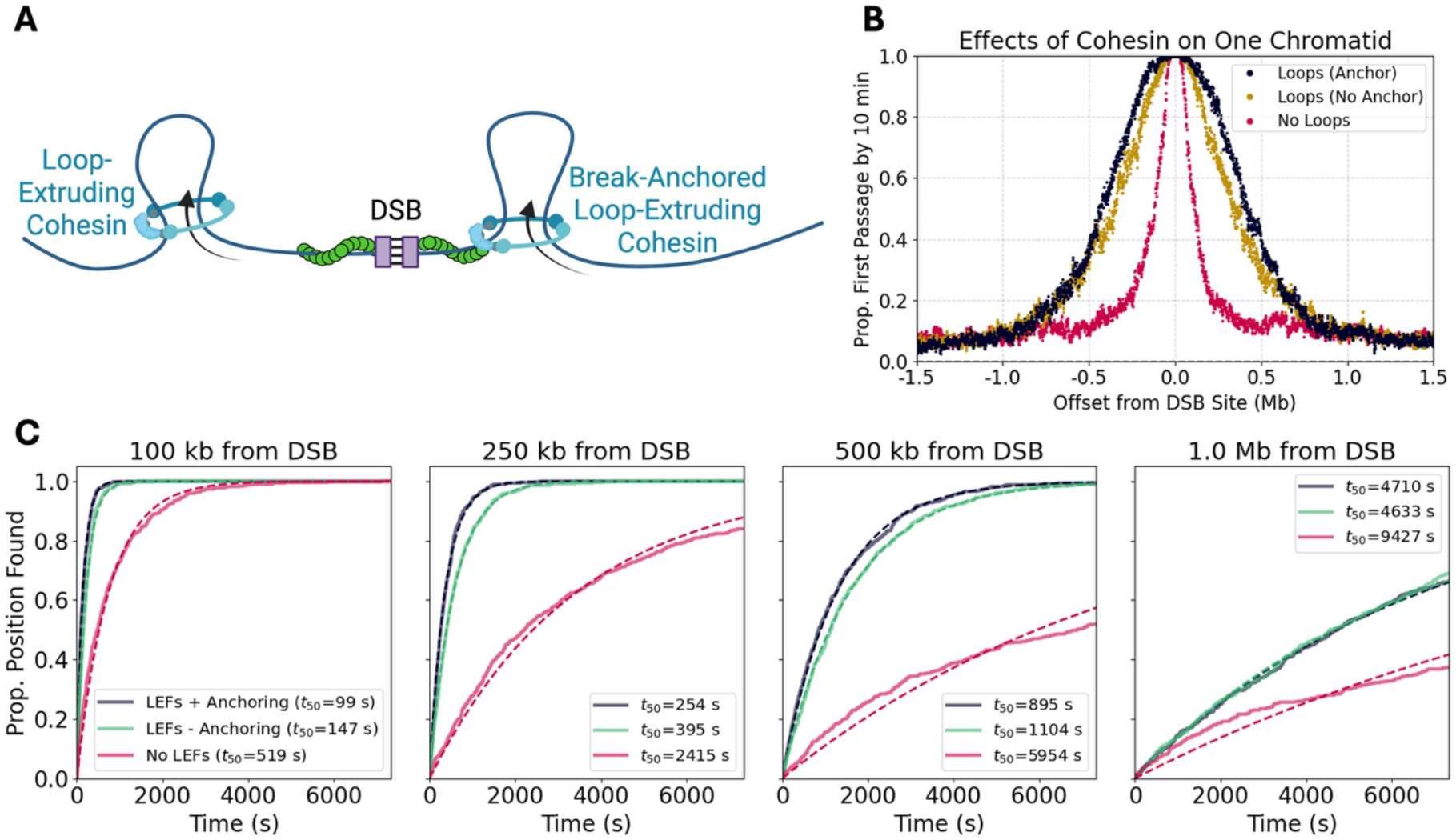
Loop-mediated chromatin reorganization on one chromatid. Simulations of one chromatid. (A) a cartoon of a DSB site (created with BioRender.com). (B) first passage attainment by 10 minutes as a function of positional offset from the DSB site. As used here, first passage attainment describes the proportion of simulations to have achieved a contact between the DSB site and a given position along the simulated polymer within a proscribed timeframe, and is a proxy for search success. (C) proportion of simulations in which positions a given distance from the DSB site has been encountered by the DSB over time, with an exponential function fitted. Positions 100 kb, 250 kb, 500 kb, and 1 Mb from the break are considered.

We first examined the single-chromatid search profile by plotting the proportion of simulation instances in which the DSB site had encountered each monomer position along the chromatid (i.e. search completion as a function of chromatin position) after 10 minutes of search time (Fig. 1B). Simulations containing loop-extruding cohesins conducted broader searches than those lacking them. Moreover, simulations in which the DSB site anchored loop-extruding cohesins showed accelerated search dynamics over those lacking anchoring for regions within 1 Mb of the break.

To explore how the search evolves over time, we selected four sample positions at 100 kb, 250 kb, 500 kb, and 1 Mb from the break, and plotted search completion to those positions as a function of time (Fig. 1C). We quantified these kinetics by fitting an exponential function to the resulting curves and obtaining *t*_50_, the time at which half of the simulations show a successful search. Anchoring cohesin at the DSB accelerated searches to donors located at 100 kb, 250 kb and 500 kb from the break by a factor of 1.65, 1.5 and 1.2, respectively; no effect on the search was visible at 1 Mb. We also tested the effects of CTCF, a transcription factor found at TAD boundaries which stalls loop-extruding cohesins, on search kinetics. While CTCF-like semi-permeable boundary elements increased search success rates at their locations, they possessed an insulating effect which hampered searches between the DSB site regions of chromatin past the CTCF, consistent with (7) (Supp. Fig. 1).

### Loop-extrusion rearranges chromatin undergoing sister chromatid searches

To investigate whether loop extrusion plays a role in mediating sister chromatid recombination we simulate the sister chromatid as a second polymer attached to the broken chromatid at randomly selected sites reflecting sister chromatid linkages. These sites correspond to TAD boundaries, consistent with (7, 19), and therefore have semi-permeable boundary element properties, enabling them to stall loop-extruding cohesins. Noting that CTCF at TAD boundaries protects loop-extruding cohesin against WAPL-mediated unloading (20, 21), we applied a fourfold increase in loop-extruding cohesin lifetime upon stalling against TAD boundaries, consistent with (11).

Following the ChIP-seq observations of (7) showing accumulation of cohesive cohesin at DSB sites, we assume that if a DSB site enters into close proximity (2 monomer-lengths) with the sister chromatid, cohesive cohesin can be recruited to the DSB and subsequently mediate cohesion between the break and the sister chromatid. The cohesive cohesins are considered bound to the DSB site, and therefore cannot translocate along the broken chromatid. However, they may translocate along the sister chromatid if they are pushed by a loop-extruding cohesin, in line with (22). We simulated 500 chromatid pairs with each chromatid spanning 2 megabase pairs (Mb), split evenly across 10 independent simulation runs. For each pair, we induced a DSB in the center of one (broken) chromatid and defined a homology site in the center of the other chromatid (the sister chromatid), to which the DSB site could bind if it entered a 2 monomer-length capture radius. A cartoon of our model is provided in Fig. 2A.

**Figure 2:**
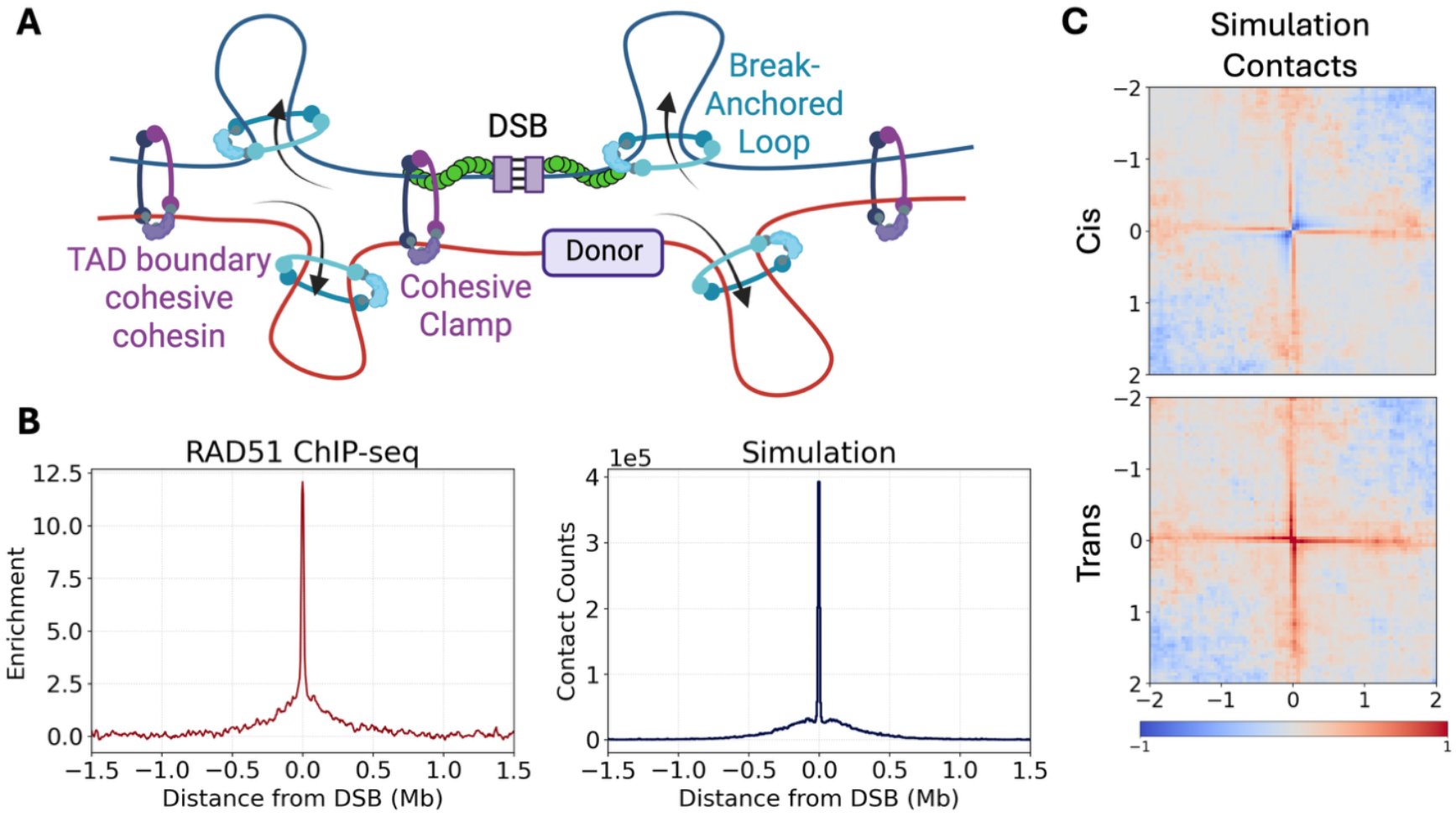
Simulating the trans search process. (A) a cartoon of the homology search process (created with BioRender.com). (B) RAD51 ChIP-seq as reported in (8) is compared to cumulative simulation contacts between the DSB site and chromatin positions. Both datasets are binned to 5 kb. (C) contact maps from homology search simulations, displayed as log_2_ (DSB/No DSB).

We first asked whether our simulations reprise the experimental homology search profiles obtained via RAD51 ChIP-Seq in (8). To do so, we recorded the simulated contact profile by analyzing DSB-chromatin interactions and applying a 5 monomer-length contact threshold. We summed DSB-chromatin contacts with the broken and sister chromatid to account for the fact that both sister chromatids are identical in sequence and therefore, undistinguishable in the ChIP-Seq experiments. We binned the resulting data and plotted each bin’s total contacts to generate a ChIP-Seq-like output. The simulation and experimental data exhibited strong agreement, with a sharp narrow peak at the DSB site and a broad peak extending 500 kb to either side of the break (Fig. 2B).

We next compared our simulations to the sister-pore-C data described in (7), which relies on BrdU labeling prior to DNA replication to distinguish intra-chromatid and inter-chromatid contacts. We generated chromosome conformation capture contact maps from our simulations, differentiating between *cis* and *trans* contacts. Our simulations successfully recapitulated contact stripes characteristic of the sister-pore-C data (Fig. 2C), with increased contacts centered at the DSB site and extending over 1 Mb in both *cis* and *trans* cases. As in sister-pore-C data, a fall-off in *cis* contacts occurs in quadrants I and III nearby the DSB site. Loop-extruding cohesins were necessary to reprise experimentally observed contact stripes and their absence produced thick, patchy *trans* contact stripes and no visible contact stripes in *cis* (Supp. Fig. 3A). Furthermore, cohesive clamps served to extend contact stripes in simulations in which loop-extruding cohesins were present, bringing those simulations into closer agreement with the sister-pore-C experiments.

These findings support that the combined presence of loop-extruding and cohesive cohesins can qualitatively capture the experimental data through their interactions with the DSB site.

### Loop-extruding cohesin decreases first passage times

To explore search kinetics, we conducted 50 simulations each containing 5 repeats of two paired chromatids of 2 Mb each, with a DSB centered along one of each of the two chromatids (totaling n=250). We then explored the profile of the homology search by plotting the proportion of simulation instances to attain first passage with (i.e. enter the proximity of) potential homology positions along the sister chromatid (Fig. 3A and Supp. Fig. 4). With a DSB site located in the center of the broken chromatid, searches to the center of the sister chromatid (directly across from the DSB site in index space) were most efficacious, with first passage attainment falling off for potential homology sites offset from the center. We compared simulations with loop-extruding cohesins and break interaction (cohesive clamps and break-anchoring) to those with loop-extruding cohesins but no break interaction, as well as to simulations with no loop-extruding cohesins. At 10, 30, and 60 minutes, loop-extruding cohesins dramatically improved search efficacy over simulations with no loop-extruding cohesins. Break interaction accelerated searches in the first half-hour of the search process, however their effects became less noticeable by 60 minutes, by which time simulations with and without break interaction had both achieved a high degree of first passage attainment.

**Figure 3:**
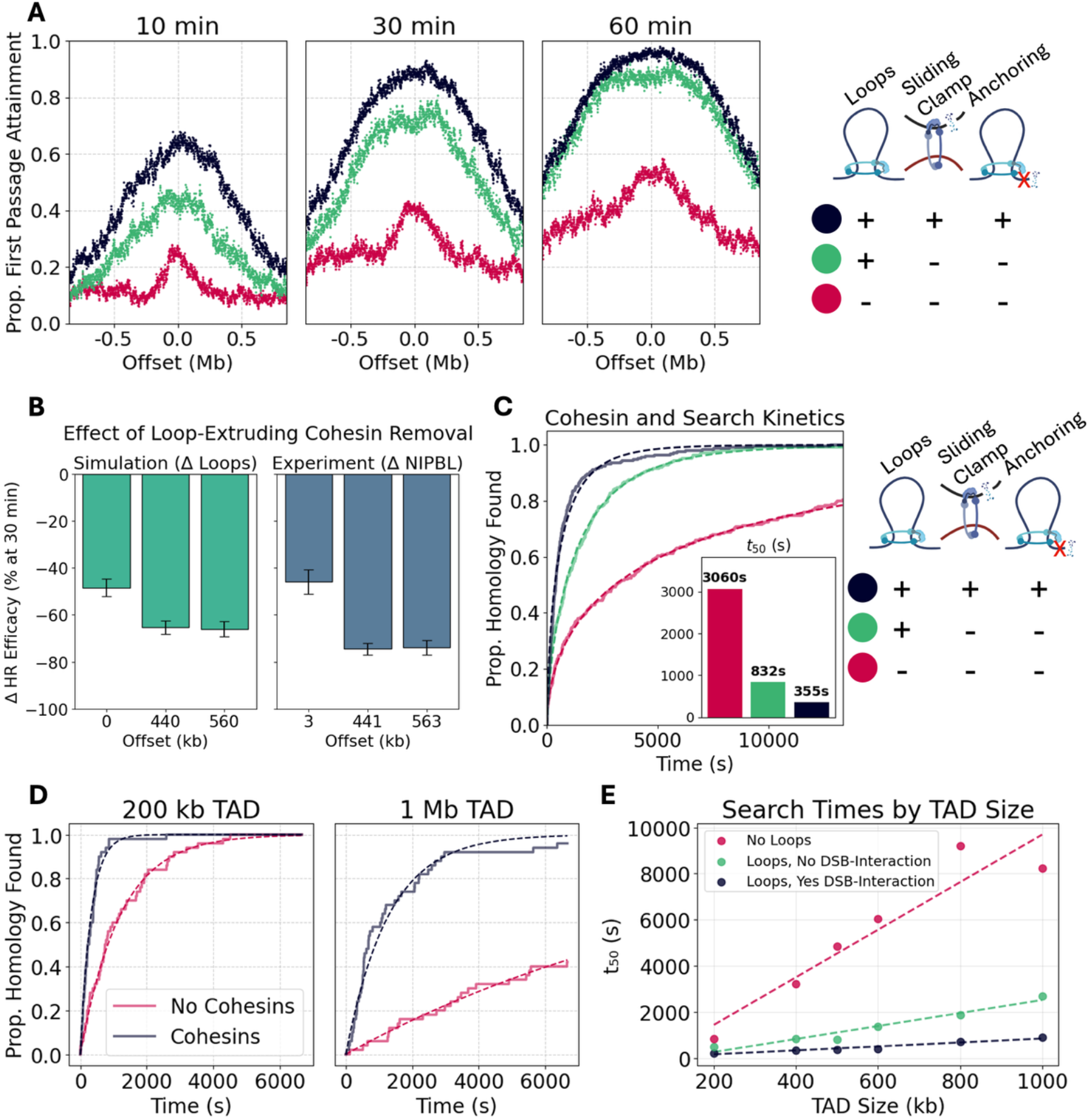
Impacts of cohesin on homology search kinetics. (A) profile of the homology search. The x-axis is sister chromatid position, measured in offset from the center of the sister chromatid (which is directly across from the DSB site in index-space, and therefore reflects the expected position of the homology site in the absence of any offset between sequence locations on sister chromatids). Proportion of simulated DSBs (out of n=250) to achieve first passage with given sites along the sister chromatid is reflected on the y-axis, serving as a measurement of homology search efficacy between the DSB site and that position. (B) comparison of simulated search efficacy to experimental homologous recombination reporter results depicting the reduction in the proportion of simulations to achieve HR success upon removal of loop-extruding cohesins. Experimental data was originally reported in (8) and used siRNA for NIIPBL depletion. Cohesive clamps may be recruited in both simulation cases. (C) proportion of simulations in which a homology site located in the center of the sister chromatid (directly “across” from the DSB site in the center of the broken chromatid) is found as a function of time, fitted to a Weibull distribution (n=250). t_50_ values obtained from this fit are provided in an adjoining bar graph. (C, D) impact of TAD size on search time to a centered homology site in simulations with fixed TAD boundaries (N=50 each). (C) search kinetics for TAD sizes of 200 kb and 1 Mb, with and without cohesin (break interaction is enabled in cases with cohesin), with exponential distributions fit. (D) right graph measures search time as a function of TAD size by using such fitted exponential distributions to solve for t_50_. The fitted lines have slopes of 10.31 (R^2^ = 0.89), 2.816 (R^2^ = 0.95), and 0.8536 (R^2^ = 0.95) for the no loops, loops without DSB-interaction, and loops with DSB-interaction cases, respectively.

We previously reported experimental measurements of HR efficiency for DSB-donor separations of 3 kb, 441 kb and 563 kb in cells with and without depletion of NIPBL, a key component of loop-extruding cohesin (8). For comparison with this experimental data, we selected positions at the center of the sister chromatid (directly “across” from the DSB, corresponding to a 0 kb DSB-homology offset) and at 440 and 560 kb from the sister chromatid center. At 10 min, 30 min and 60 min timepoints, we compared the relative search efficiency between simulations with and without loop-extruding cohesins, holding all other parameters (including cohesive clamp recruitment) constant. At 30 min, we found an approximately 50% reduction in search success at 0 kb offset upon loop-extruding cohesin removal, and an approximately 70% reduction in success at 440 and 560 kb. Assuming that a single DSB-donor encounter results in successful HR, these findings align well with experimental measurements of HR efficiency upon depletion of NIPBL (Fig. 2B). Qualitatively similar results were obtained for the 10- and 60-minute timepoints, although the experimental agreement was lower for these timepoints than for the 30 min one (Supp. Fig. 3C).

To understand how interactions between DSB sites and cohesin influence search outcomes we considered scenarios with different cohesin properties. We kept break-independent loop-extruding cohesin activity constant while disabling the DSB-associated cohesive clamp, break-anchoring, or both (Supp. Fig. 3D). We found double-digit percent reductions in search efficacy in all three cases, suggesting that interactions between DSB sites and cohesin contribute to ensuring successful homology searches and, ultimately, timely repair.

### Effect of TAD size on cohesin-mediated searches

To uncover search kinetics, we plotted the proportion of simulation instances (fraction of DSB sites) to contact a homology position centered on the sister chromatid over time (Fig. 3B). Simulations with loop-extruding cohesins found the homology site much more rapidly than those mediated by passive diffusion alone. Simulations with both loop-extruding cohesin and DSB-interactions (defined as cohesive clamp recruitment and break-anchoring) were faster yet. We determined that our data fit a Weibull cumulative distribution function, mathematically described as 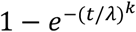 where *t* is time and λ and *k* are free parameters. We used this fit to obtain *t*_50_ (time at which half of all simulations have found the homology site) values for each case, finding a *t*_50_ of 3060 seconds for passive-diffusion-only searches, 832 seconds upon loop-extruding cohesin introduction, and 355 seconds when loop-extruding cohesins were present and DSB-interactions were enabled.

Our fitted distributions involved values for *k* of 0.54 for simulations without loop-extruding cohesins and 0.73 and 0.69 for simulations with loop-extruding cohesins without and with DSB-interaction, respectively. These values relate to the time-dependent hazard rate of our simulations. In a Weibull model, values of *k* < 1 indicates a decreasing hazard over time, implying that homology encounters are more likely at early timepoints in the simulation and become progressively less likely at later times. We posited that this phenomenon stemmed from our simulation conditions, in which TAD sizes were randomized. We therefore simulated n=50 DSB instances set in pairs of 1.4 Mb chromatids with DSBs centered in pre-defined TAD boundaries, for TAD boundaries of 200 kb to 1 Mb in size. We found that search attainment over time to homology positions centered in the sister chromatid fit to an exponential distribution in these simulations, and by plotting *t*_50_ as a function of TAD size we determined that larger TADs correlate with longer search times (Fig. 3C). Moreover, we found that the effect of TAD size on search times, quantified using the slope of a fitted line, was greater for passive-diffusion simulations with no loop-extruding cohesins (10.31 s/kb) than for simulations with loop-extruding cohesins (2.816 s/kb without break interaction), and greater for simulations with loop-extruding cohesins but lacking break interaction than for those with break interaction (0.8536 s/kb). Thus, both DSB-independent cohesins and DSB-interacting cohesin accelerate search dynamics, and this effect is more pronounced as TAD size increases. These results suggest that cohesin-driven homology searches are particularly helpful for DSBs that occur in large TADs.

### A cohesin-driven mechanism for homology search

Previous works have shown increased recruitment of loop-extruding cohesins to chromatin regions undergoing homologous recombination (7, 8), however it is unclear whether this break-associated increase in cohesin density favors one chromatid or is evenly distributed across both chromatids during repair. We thus sought to address the relative contributions to the search process from loop-extruding cohesins on the broken chromatid and from those on the sister chromatid.

We designed simulations with loop-extruding cohesins present on the broken chromatid or sister chromatid alone and compared their search kinetics with simulations with loop-extruding cohesins on both chromatids and with simulations lacking loop-extruding cohesins (Fig. 4A). We enabled cohesive clamp recruitment to the break site in these simulations. As expected, the presence of loop-extruding cohesins on both chromatids resulted in a significant reduction of search times compared to the no loop-extruding cohesins case (*t*_50_ = 357 for loop-extruding cohesins vs *t*_50_ = 2318 s for no loop-extruding cohesins). Notably, simulations with only broken chromatid loop-extruding cohesins exhibited similar search kinetics ( *t*_50_ = 2614 s) to those with no loop-extruding cohesins at all. In contrast, simulations with loop-extruding cohesins on the sister chromatid yielded a search time of *t*_50_ = 783 s, somewhat slower than the both chromatids case, but substantially faster than the no loop-extruding cohesins one. These results suggest that loop-extrusion along the sister chromatid is the major contributor to the cohesin-driven search. Interestingly, although loop-extrusion on the broken chromatid alone does not improve search kinetics, it accelerates searches when working in synergy with sister chromatid extrusion.

**Figure 4:**
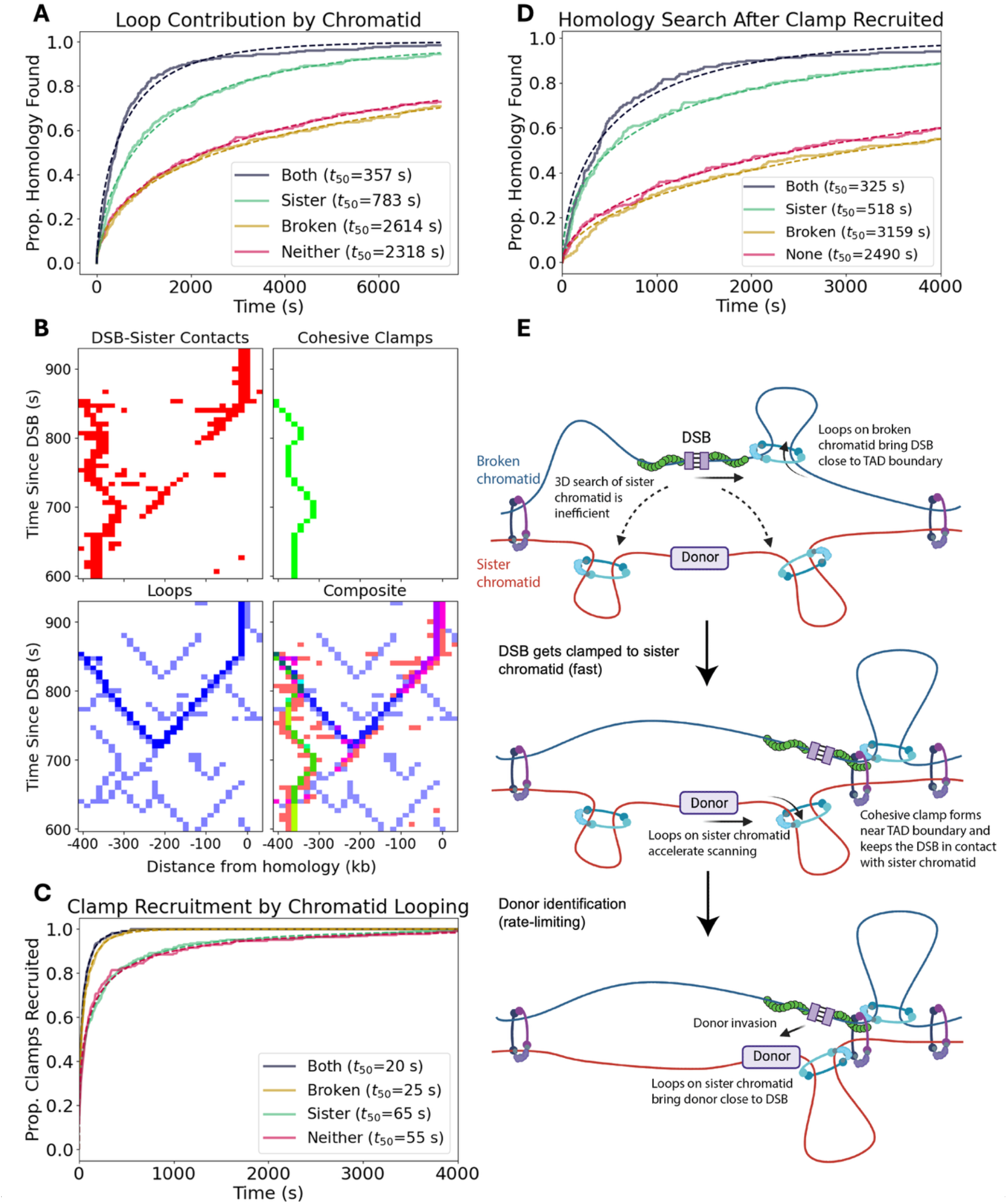
Chromatid-specific effects of loop-extruding cohesin. (A) search kinetics following DSB induction for simulations with loop-extruding cohesin present on only one chromatid are compared with those of simulations with loop-extruding cohesin present on both chromatids and simulations allowing cohesive clamp recruitment but lacking loop-extruding cohesins. (B) a representative contacts trace of a single DSB instance with a sliding clamp cohesive cohesin bound. Contacts (threshold = 5 monomer-lengths) between the DSB site and positions along the sister chromatid are displayed with respect to time in red. Sliding clamp cohesive cohesin positions along the sister chromatid are shown in green, while sister chromatid loop-extruding factors are portrayed in blue. A darker shade of blue is used to highlight the loop-extruding factor which was proximal to (threshold = 10 monomer-lengths) the homology site at the time the DSB first contacted and bound the homology. TAD boundaries are present at 500 kb to either side of the homology site. (C) kinetics of cohesive clamp recruitment, where the x-axis represents time since clamp recruitment was first enabled in the simulation following DSB induction. (D) search time between clamp recruitment and homology identification. (E) a cartoon of the homology search process (created with BioRender.com). All t_50_ values determined using a Weibull fit.

To better understand the mechanism by which loop-extruding cohesins reduce search times, we turned to single-break traces. We visually inspected 50 traces from DSB sites centered within 1 Mb TADs and determined that of the 41 traces in which the DSB found the homology site within the simulation timeframe (5000 timesteps, or 3,333 seconds), approximately two thirds (n=27) accomplished this in two stages. First, the DSB contacts the sister chromatid, allowing a cohesive clamp to load and stabilize the inter-chromatid contact. Then, loop-extruding cohesins reel in the sister chromatid, presenting it to the DSB (Fig. 4B).

We studied the kinetics of these two stages – initial contact with the sister chromatid and subsequent 1D scanning – in simulations with and without loop-extruding cohesins. We found that both stages were significantly accelerated by chromatin loops. Loop-extruding cohesins accelerated clamp recruitment from *t*_50_ = 55 s in the absence of looping to *t*_50_ = 20 s when loop-extruding cohesins were present, thus reducing the time for initial contact between the DSB site and the sister chromatid (Fig. 4C). We next calculated times elapsed between clamp binding and DSB-homology contact. Again, loop-extruding cohesins caused a dramatic acceleration, from *t*_50_ = 2490 s without loop-extruding cohesins to *t*_50_ = 325 s in their presence (Fig. 4D). Notably, clamp recruitment occurred on much faster timescales than post-clamp searches (Supp. Fig. 4D), suggesting that the latter is the homology search’s rate-limiting step.

We sought to determine what roles broken and sister chromatid loop-extruding cohesins played during these two stages. We found that sister chromatid loop-extruding cohesins alone did not accelerate clamp recruitment beyond the no loop-extruding cohesins case. Broken chromatid loop-extruding cohesins, however, were both necessary and sufficient to restore *t*_50_ values to close to those for simulations with loop-extruding cohesins on both strands ( *t*_50_ = 25 s with broken chromatid loop-extruding cohesins only), suggesting that they are the primary driver of initial contact between the DSB and the sister chromatid. This may be achieved by bringing the DSB close to the TAD boundary, where both sisters are physically connected. Indeed, clamp recruitment happened predominantly at TAD-proximal sites, and this trend was abolished in simulations lacking broken chromatid loops (Supp. Figs. 5A and 5B). Conversely, broken chromatid loop-extruding cohesins alone had no impact on post-clamp searches, which were accelerated to *t*_50_ = 518 s in simulations with only sister chromatid loop-extruding cohesins (Fig. 4D). This indicates that sister chromatid extrusion is the primary driver of post-clamp search processes.

These findings suggest a role for broken chromatid loop-extruding cohesins in facilitating initial contact between the DSB site and the sister chromatid – possibly by bringing the DSB close to a TAD boundary – followed by cohesive-cohesin-mediated stabilization of DSB-sister chromatid contacts (Fig. 4E). Sister chromatid loop-extruding cohesins may then drive the search process after the DSB is clamped by sequentially presenting chromatin to the break via loop extrusion. This model provides an explanation for why sister chromatid loops are the dominant contributor to search process: these loops drive scanning along the sister chromatid, which is the rate-limiting step.

## DISCUSSION

Recent works have shown that loop-extruding cohesin plays a key role in the genomic searches that accompany homologous recombination repair (7, 8), however the underlying mechanism remained unexplored. Our simulations recapitulate several experimental findings (RAD51 ChIP-Seq, sister-pore-C, and HR-GFP assays) and suggest a model for cohesin-driven search supporting sister chromatid recombination. Notably, our computational findings indicate that loop extrusion alone does not achieve optimal donor identification. Instead, repair is greatly accelerated by interactions between the DSB site and cohesin, including break-anchoring and cohesive cohesin recruitment. Moreover, we find that searches are dominated by loop-extruding cohesin on the sister chromatid, with a supporting role played by loop extrusion on the broken chromatid.

Based on our findings, we propose that the homology search occurs in a two-stage process. In the first stage, broken chromatid loop-extruding cohesins bring DSB sites close to TAD boundaries (Supp. Figs. 5A and 5B), facilitating DSB-sister chromatid contacts and cohesive clamp recruitment (Fig. 4C). Subsequently, sister chromatid loop-extruding cohesins scan the DSB along the sister chromatid to accelerate identification of the homology site (Fig. 4B). This model converts the search for a donor template on the sister chromatid from passive 3D diffusion to cohesin-driven 1D scanning.

These two stages – cohesive clamp recruitment and search along the sister chromatid – are not independent of each other. For instance, sister chromatid scanning may benefit from the positioning of cohesive clamps near TAD boundaries, sites of intense loop-extruding activity that could further enhance the search (Supp. Figs. 5C and 5D). In fact, we analyzed single-break traces in which cohesive clamp recruitment was disabled and found that, in many instances, looping DSB sites to TAD boundaries facilitated scanning by loop extrusion on the sister chromatid (Supp. Fig. 6). We also note that while our simulations assume *de novo* cohesive clamp recruitment, it may be possible for loop-extruding cohesins to translocate cohesive cohesin from elsewhere on the genome to break sites, similarly to how cohesive cohesin is likely positioned at TAD boundaries (22). In such an event, broken chromatid loop-extruding cohesins would play an additional role in establishing a 1D search, as they would drive cohesive cohesin translocation onto break sites.

Future works may seek to explore the mechanisms underlying elevated loop-extruding cohesin recruitment observed at DSB sites (7, 8), which could lead to new lines of inquiry into cohesin’s effects on the search process. Additionally, the mechanism by which cohesin is anchored at the DSB has not yet been established, and could merit structural and kinetic studies. Cohesin is a ubiquitous regulator of genome architecture, and our data show how it can dramatically accelerate genomic searches within and between replicated chromatids by combining its cohesive and loop-extruding activities.

## METHODS

### Loop-extruding factor translocation

LEF translocations are calculated in one dimension as in (9). Each LEF has two legs, which occupy sites along a lattice representing a chromatid. Every loop extrusion timestep, each leg may translocate a single step away from the other leg along the lattice. Legs may stall at boundary elements with a b=0.5 probability and will also stall upon encountering other loop-extruding cohesins. The DSB itself has a 100% probability of stalling loop-extruding cohesins. Like loop-extruding cohesins, cohesive clamps recruited to DSB sites have two legs, with one leg on each chromatid. The DSB-bound leg of the cohesive clamp acts as a boundary element, while its sister chromatid leg may be freely pushed by loop-extruding cohesins without impeding their movement.

Loop-extruding cohesins are subject to processivity *λ*=120, defined as lifetime multiplied by the extrusion rate. The processivity of loop-extruding cohesins may be modified upon capture by a boundary element. They also have an initial average separation *d*=120, with the total number of loop-extruding cohesins present in the simulation given as the number of lattice sites divided by *d*. These values are consistent with experimental and computational literature (9, 18). Every loop extrusion timestep, each LEF has a probability to unload given by (λ/extrusion rate)^−1^. When a LEF unloads, another is immediately loaded at a random unoccupied site on the lattice to maintain the same density of loop-extruding cohesins. Consistent with previous works showing increases in cohesin lifetime when stalled at CTCFs (11), a fourfold increase in processivity was applied to LEFs stalled at DSB sites or at boundary elements representing CTCFs, including TAD boundaries.

### 3D molecular dynamics

The Polychrom library (23), a wrapper for OpenMM (24), is used to conduct 3D molecular dynamics simulations. Here, chromatin is represented as a polymer, in which each monomer represents 1 kb of chromatin and corresponds to an LEF translocation lattice site. To convert simulation timesteps to seconds, we assume a loop extrusion rate of 3 kb/s, consistent with (25). Monomers are connected to each other by harmonic bonds, described by 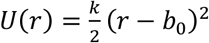 where *r* is the distance between monomers, *b* is the diameter of a monomer, and *k* = 2*k*_*B*_*T*/*δ*^2^ where *k*_*B*_ is the Boltzmann constant, *T* is the temperature, and *δ* is 0.05 monomers, representing the average fluctuation in bond distance. Identical bonds link monomers whose 1D lattice sites are occupied by the same loop extruding factor. Such harmonic bonds are also used to represent cohesive cohesins in two-strand simulations. To prevent chromatin from crossing itself, a polynomial repulsive potential is applied between monomers, given by 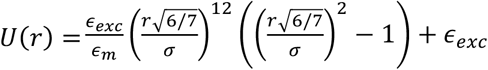 for *r* < σ = 1 monomer, where *ϵ*_*m*_ = 46656/823543 and *ϵ*_*exc*_ = 3*k*_*B*_*T*. An angular force of 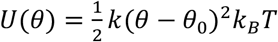 with *k* = 1.5 and θ_0_ = π is also applied. 750 molecular dynamics steps were conducted for each loop extrusion timestep. Monomer positions in 3D space are recorded every 10 loop extrusion timesteps.

Homology search simulations contain two paired polymers, each representing one chromatid. Unless otherwise specified, TAD boundary positions are randomized, with an average of one TAD boundary every 500 kb. Starting from 100 timesteps after DSB induction, sliding clamp cohesive cohesins may be recruited at either side of the DSB with a programmable 4 kb offset, consistent with (7). These sliding clamp cohesive cohesins will be recruited in cases where the DSB-adjacent site is within 2 monomer-lengths of a position along the sister chromatid. Likewise, in contact profiling (ChIP-seq comparison, chromosome conformation capture, and single-break traces) simulations, the DSB may bind to pre-defined homology sites along the sister chromatid if it is within 2 monomer-lengths of that site, in which case a new bond is introduced between the homology and DSB site. The homology site corresponding to the left (lower-indexed) side of the DSB is offset from that corresponding to the right (higher-indexed) side of the DSB by a single index value.

As an equilibration step, 10,000 1D loop extrusion steps were conducted prior to the beginning of the 3D molecular dynamics simulations. An additional 1,000 loop extrusion steps (and 750,000 associated molecular dynamics timesteps) were conducted in the pre-DSB regime prior to DSB induction.

### Single-chromatid searches

Single-chromatid search simulations employ a continuous polymer composed of 150,000 monomers representing 150 Mb of chromatin. 50 DSBs are induced at evenly spaced intervals along this polymer by giving impermeable boundary element properties to the monomer positions flanking the break. A contact radius of 5 monomer-lengths was used to identify passages between the DSB site and other monomers, consistent with (9, 14, 26). In kinetics plots exploring searches to positions a fixed distance from the break, positions both that distance upstream and downstream of the break were considered to achieve better averaging. TAD boundaries and cohesive clamps are not present in all single-chromatid simulations, however the effects of TAD boundary CTCFs are explored in Supp. Fig. 1, where boundary elements representing CTCFs are present at set locations along the chromatid.

### DSB search contact profiling

Contact maps and ChIP-seq comparison contact profiles were generated using a cutoff radius of 5 monomer-lengths. 5000 post-DSB loop extrusion steps were conducted for each simulation. Across 10 simulation runs of 50 DSB instances each, this leads to a total of 250,000 polymer conformations arising from n=500 independent DSB instances. Single-break traces were similarly obtained by applying a contact radius of 5 monomer-lengths at each recorded timepoint to individual simulation runs containing n=50 DSB sites with 1.4 Mb of chromatin per chromatid and fixed TAD boundary positions.

### Search kinetics simulations

Simulations exploring homology search kinetics and cohesive clamp recruitment applied a 5 monomer-length cutoff radius to determine whether DSB sites had achieved passage with potential homology sites. Passages were tallied starting from 30 loop extrusion steps after the timepoint of DSB induction. DSBs do not bind defined homology sites in these simulations, allowing us to profile searches to positions along the entire sister chromatid. Unless otherwise stated, data in these simulations were drawn from 50 independent simulation runs containing 5 sister chromatid pairs each with 2 Mb of chromatin per chromatid (n=250 DSB sites). In simulations with chromatid-specific LEF recruitment, LEF densities were held constant at *d*=120 by halving the total number of loop-extruding cohesins in the simulation compared to simulations in which loop-extruding cohesins may be recruited to either chromatid.

## CODE AVAILABILITY

Simulation and analysis code used in this work will be made available on GitHub upon publication.

## SUPPLEMENTARY FIGURES

**Supplementary Figure 1:**
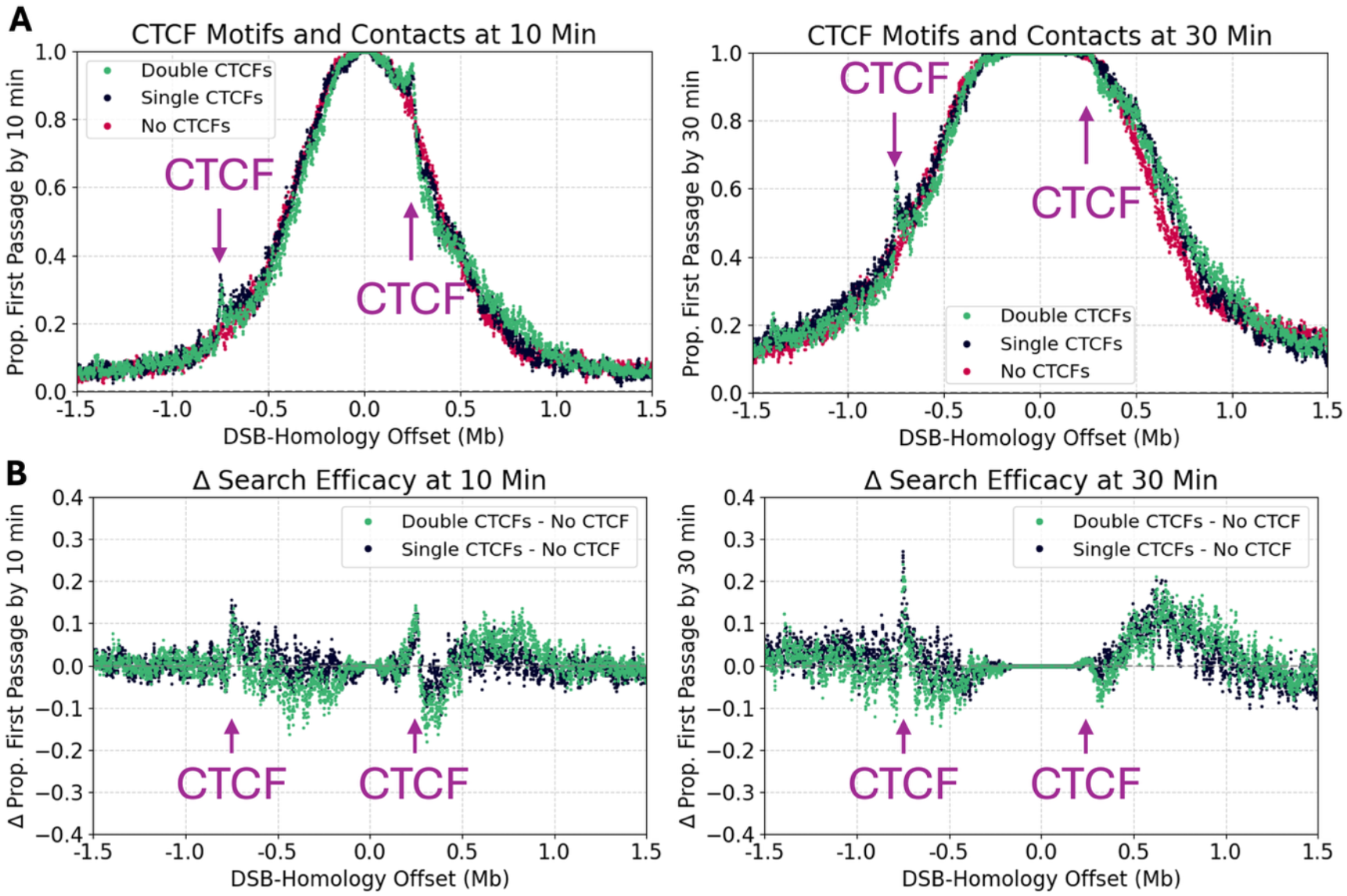
CTCF and single-chromatid searches. Single-chromatid simulations of DSB contact profiles on chromatids including boundary elements representing CTCF at -750 kb and +250 kb offsets from the break site. Positions of CTCFs are indicated by purple arrows. (A) effects of a single CTCF motif and of a double CTCF motif on first passage attainment profiles, in which two CTCFs are bound at a 10 kb separation from each other (-5 kb and +5kb of the original location). (B) presents the difference in first passage attainment between simulations including and not including CTCF motifs as a function of distance from the DSB site.

**Supplementary Figure 2:**
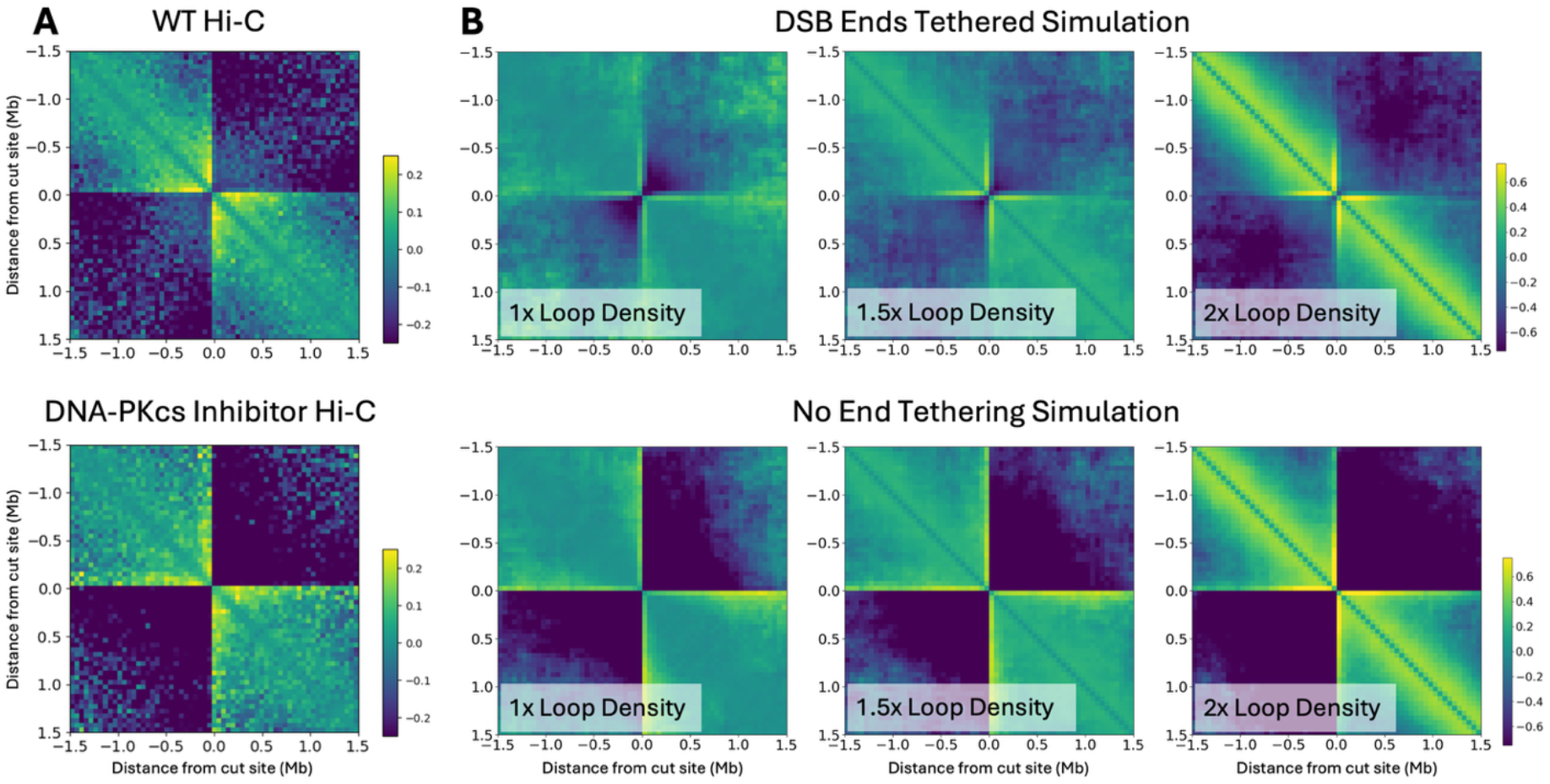
Contact maps of single-chromatid searches. (A) experimental Hi-C data from wild-type cells and cells with DNA-PKcs inhibited, previously reported in (8). PNA-PKcs is a protein associated with end tethering in NHEJ repair and is an early responder to DSBs. (B) simulation single-chromatid contact maps with and without DSB end tethering and with varying extents of post-break loop-extruding cohesin recruitment. End tethering was disabled by cleaving the bond between DSB ends, allowing them to diffuse freely in space. Loop-extruding cohesins were recruited immediately after the break to the specified total density. Simulations are across n=500 instances, distributed across 10 runs each containing a 150 Mb polymer segmented into 50 regions of 3 Mb each, with a DSB centered in each region.

**Supplementary Figure 3:**
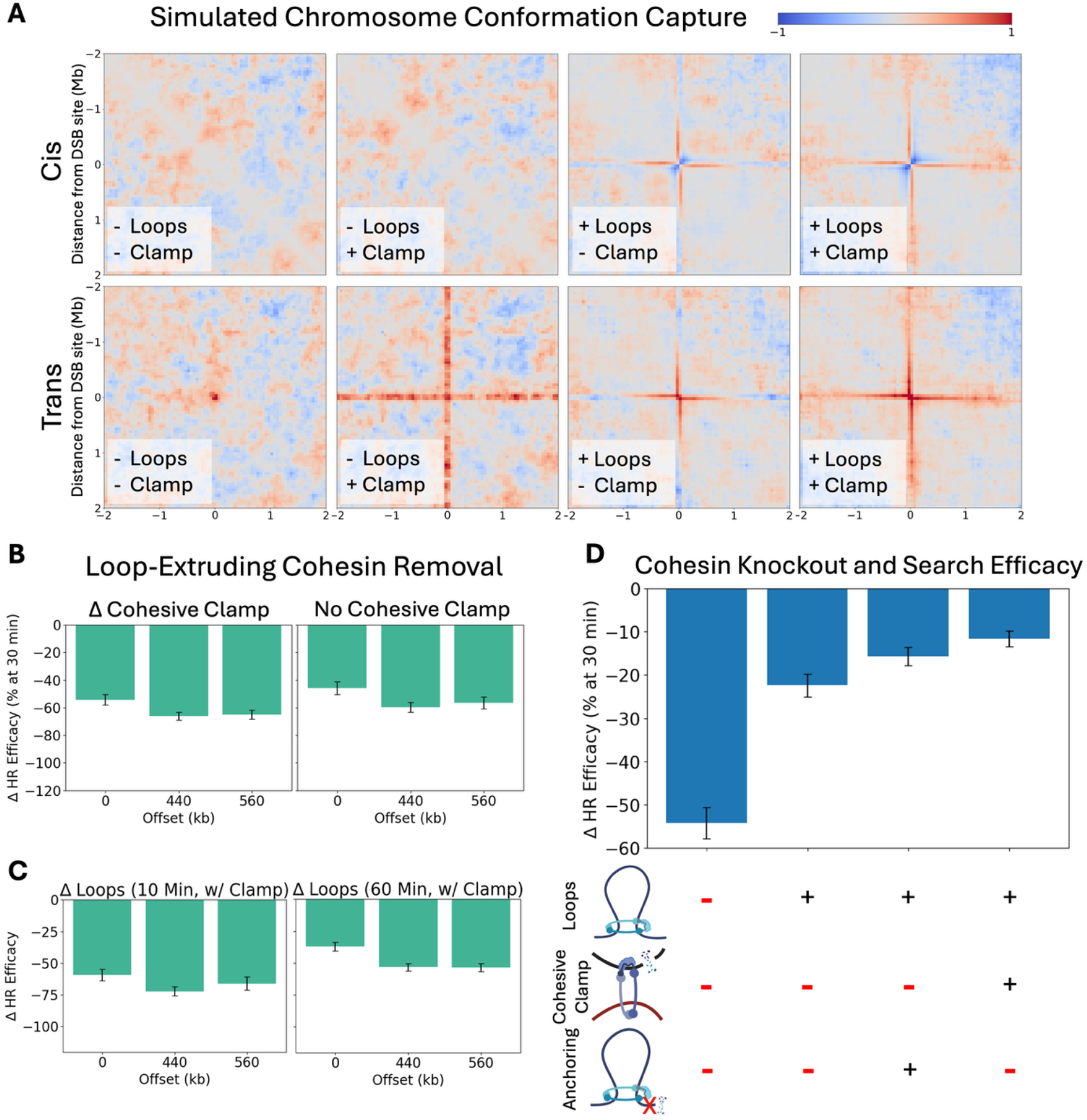
Homology search contact maps and HR efficacy. (A) contact maps from two-chromatid simulations with and without loop-extruding cohesins and cohesive clamps. (B) change in HR efficacy (first passage attainment) in simulations upon loop-extruding cohesin removal, with cohesive clamps either removed simultaneously with loop-extruding cohesins (left) or absent from both the control and the loop-extruding cohesin removal case (right). (C) change in HR efficacy at the 10- and 60-minute timepoints. (D) change in HR efficacy from removal of different combinations of loops, clamps, and anchoring from searches to the center of the sister chromatid (0 kb offset).

**Supplementary Figure 4:**
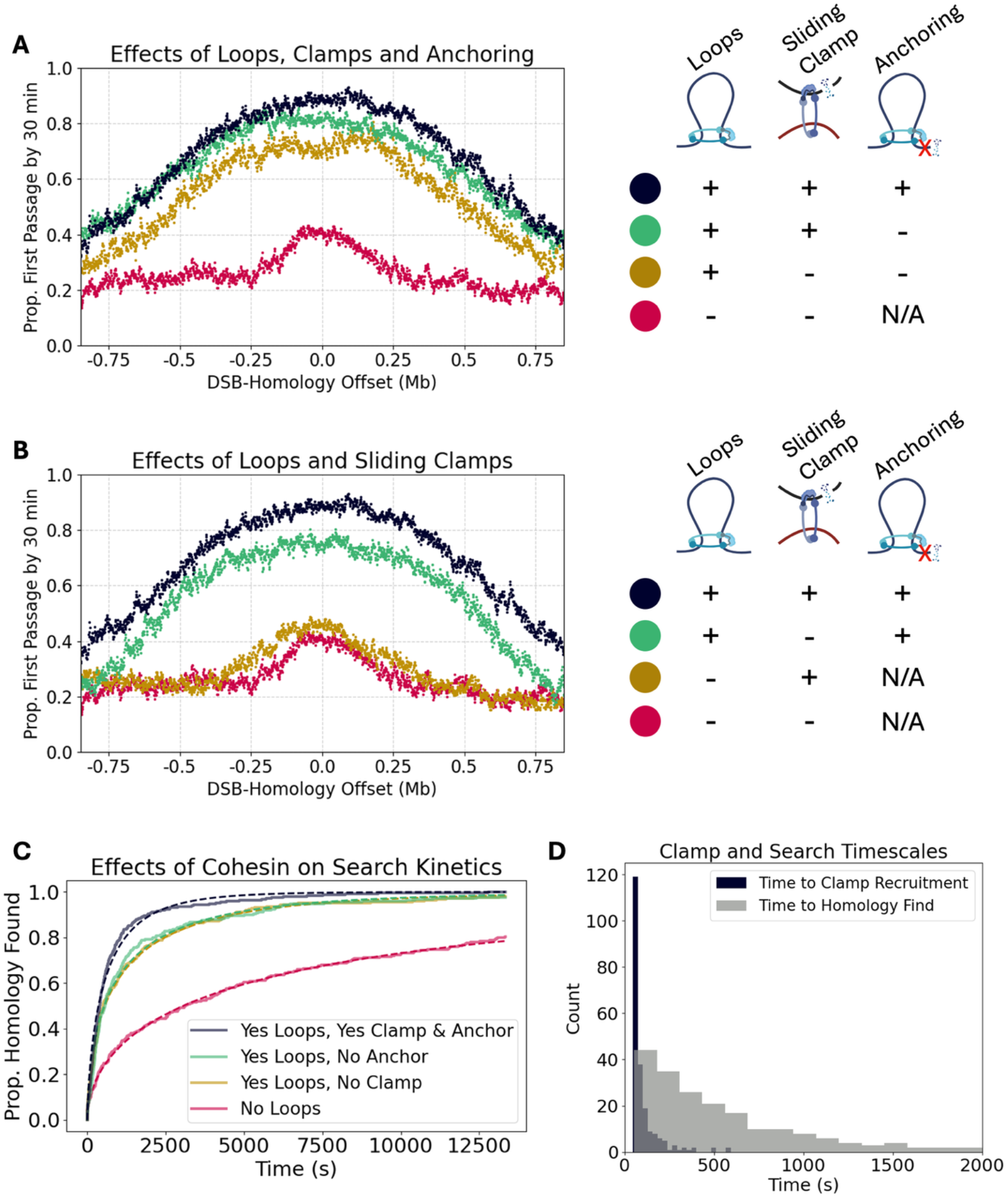
Effects of various forms of cohesin on the homology search. (A, B) homology search profile along the sister chromatid by position offset from the center of the sister chromatid, with various combinations of loops, cohesive clamps, and break-anchoring enabled. (C) effects of clamps and anchoring on search kinetics. (D) histogram of cohesive clamp binding time and homology passage time across n=218 simulation instances for a homology site centered in the sister chromatid. Simulation instances were drawn from an initial population of n=250, which were filtered to select simulations involving clamp binding prior to homology identification.

**Supplementary Figure 5:**
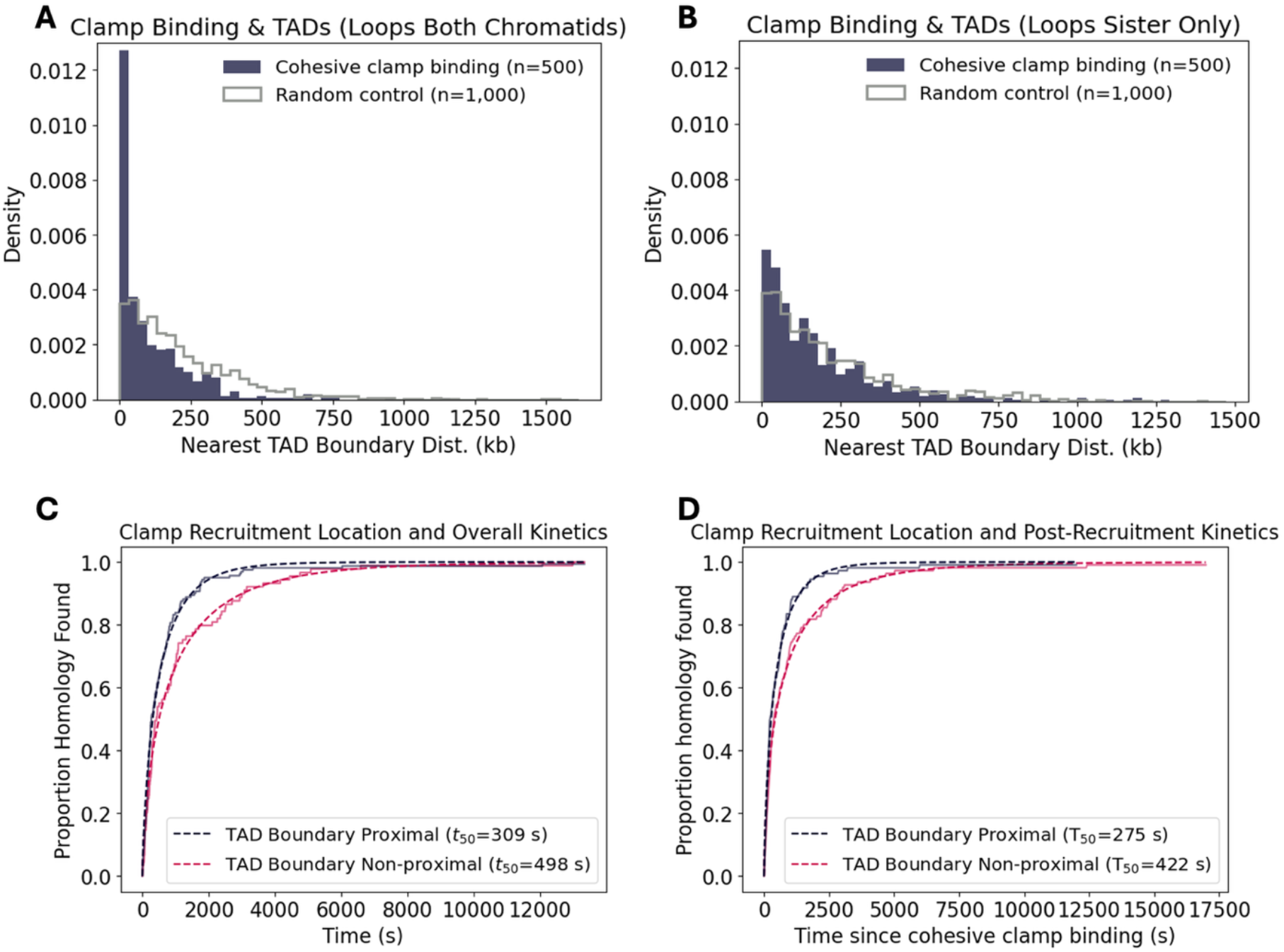
Cohesive clamp recruitment and TAD boundary locations. (C) a histogram of distances between template chromatid binding sites of DSB-recruited cohesive cohesins and their nearest TAD boundaries, with distances between TAD boundaries and randomly selected control positions in gray. (D) same histogram plotting method as (C) using data from a simulation with loop-extruding factors on the sister chromatid only. (C, D) search kinetics are graphed using a proximity threshold of 50 kb to classify DSB sites based on whether cohesive clamp recruitment events were proximal or non-proximal to TAD boundaries. (C) considers time between DSB induction and homology passage, while (D) considers time between clamp recruitment and homology passage. Homologies are assumed to be in the center of the sister chromatid. Simulation instances were drawn from an initial population of n=250, which were filtered to select simulations involving clamp binding prior to homology identification.

**Supplementary Figure 6:**
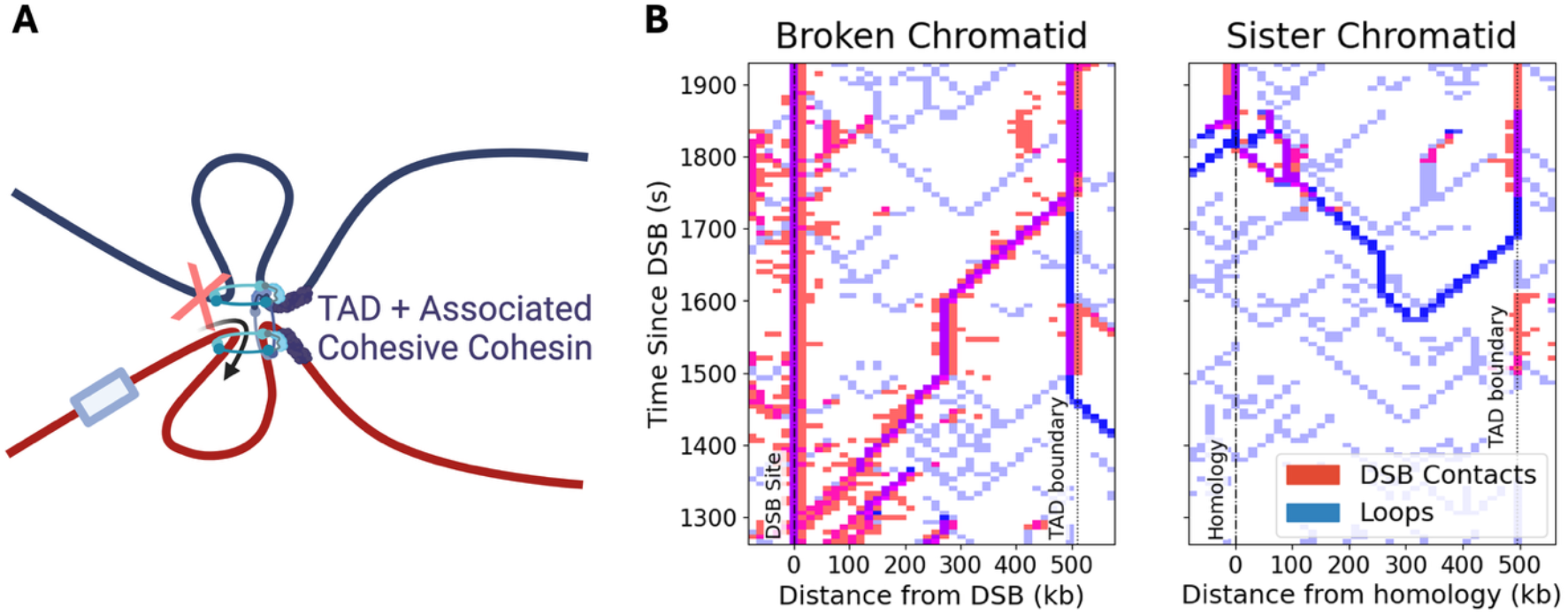
Clamp-independent homology search mechanism. (A) a model of a TAD-boundary-mediated sister chromatid search, where a broken chromatid loop-extruding cohesin loops the DSB site to a TAD boundary, enabling loop-extruding cohesins stalled at the TAD boundary on the sister chromatid to scan the DSB site along the chromatid. (B) a representative trace from a simulation with fixed TAD boundaries 500 kb from the break site (1 Mb TAD) and no cohesive clamp recruitment, showing such a mechanism.

**Supplementary Figure 7:**
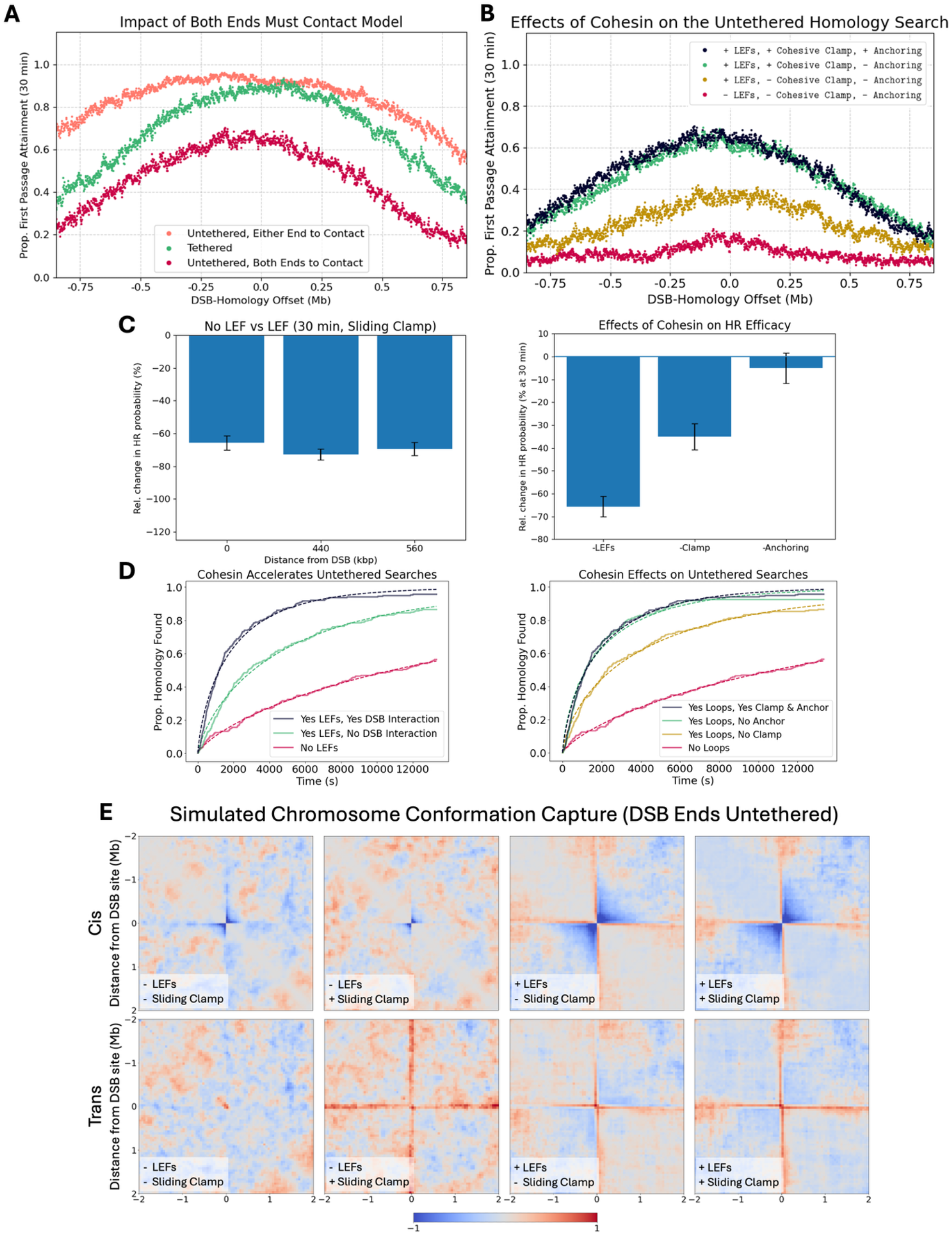
The untethered search. (A) impact of different models of loss of end tethering on the sister chromatid homology search profile. In the “both ends to contact” model, homology passage requires both DSB ends to have contacted the homology site, while “either end to contact” simply requires either end to have achieved passage. Where relevant, analysis in this figure requires both ends to have found the homology site to constitute a successful homology search. (B) effects of cohesin on the profile of the untethered homology search, showing a strong effect for loop-extruding cohesins and for cohesive clamps, but not for anchoring. (C) relative change in passage attainment for untethered searches upon loop-extruding cohesin removal. Various offsets between the sister chromatid center and the homology site are considered (left), as are the effects of cohesive clamp or anchoring removal on searches to the center of the sister chromatid (right), showing similar results to tethered searches except that break-anchoring no longer appears to improve search outcomes. (D) kinetics of untethered searches. (E) contact maps for untethered searches. n=250 for (A, B, C, D), n=500 for (E). “LEF” refers to loop-extruding factor, a term which describes loop-extruding cohesins. Loss of break-anchoring in untethered searches is here defined by unloading of loop-extruding cohesins at the break site (“falling off”).

